# Multimodal Connectivity-based Cortical Segmentation with Graph Neural Networks

**DOI:** 10.1101/2025.10.22.683959

**Authors:** Agata Łabiak, Anees Kazi, Chantal Pellegrini, Aina Frau-Pascual, Iman Aganj

## Abstract

Due to the significant amount of time and expertise needed for manual segmentation of the brain cortex from magnetic resonance imaging (MRI) data, there is a substantial need for efficient and accurate algorithms to replace the need for human involvement. In this work, we explore the capabilities of Graph Neural Networks (GNNs) to segment the brain surface based on structural brain connectivity. We train three different GNN architectures, the Graph Convolutional Network (GCN), the Graph Attention Network (GAT), and the Graph U-Net, and evaluate their performances when trained on silver-standard cortical region labels created by FreeSurfer. We take a multimodal approach to brain segmentation by examining the influence of the structural connectivity values inferred from diffusion MRI (dMRI) in addition to using values from structural MRI (sMRI). Our results demonstrate the utility of GNN models, particularly the GAT architecture, which achieved Dice scores competitive to those reported in the literature with non-graph methods. Additionally, structural connectivity derived from dMRI revealed significant value in improving automatic segmentation, as models trained on combined attributes from dMRI and sMRI outperformed those trained only on sMRI. Finally, we compared the GNN-based and the FreeSurfer segmentations in their ability to predict demographic/clinical data, where neither of the two approaches was statistically significantly superior to the other.

## 1 INTRODUCTION

Development of *in vivo* imaging techniques such as Magnetic Resonance Imaging (MRI) has given neuroscientists an unprecedentedly detailed and non-invasive view of the human brain. However, as the brain is a highly complex structure consisting of billions of neurons, interpreting MRI images remains a challenge. Brain segmentation provides a valuable way of reducing dimensionality by dividing the brain into a limited number of regions of interest (Gallardo Diez, 2018; Wang et al., 2018; Desikan et al., 2006). Further, segmentation greatly aids medicine and neuroscience by helping to advance our understanding of neuroanatomy and providing deeper insight into the connection between brain structure and human clinical and behavioral data (Gallardo Diez, 2018).

Manual brain labeling is a tedious and time-consuming process (Roy et al., 2018). As a result, there has been substantial research to fully automate the process. The vast majority of published methods use only structural MRI (sMRI) images as input, which provide important information about brain anatomy and allow measurements such as cortical thickness and curvature (Roy et al., 2018; Henschel et al., 2020, 2022; Fischl et al., 2004; Zhang et al., 2024). However, recent studies indicate that multimodal segmentation can provide novel insights and improve the utility of surface segmentation (Wang et al., 2018; Zhang et al., 2024). Specifically, the relationship between structural connectivity and cortical morphology has been exploited to leverage structural connectivity to segment (Zhang et al., 2024; Eschenburg et al., 2021; Wang et al., 2018) and align (Petrović et al., 2009; Isaev et al., 2017; Cole et al., 2025) the cortex. Functional connectivity obtained from functional MRI has also been used for such purposes (Wang et al., 2018; Conroy et al., 2013). We explore here the impact of including structural connectivity, which is a measure of the strength of physical white-matter connections between brain regions, to segment the cortex.

We train Graph Neural Networks (GNNs), such as the Graph Convolutional Network (GCN), the Graph Attention Network (GAT), and the Graph U-Net architectures, based on information from sMRI alone as well as combined sMRI and dMRI connectivity features, using the segmentation results obtained from FreeSurfer (Fischl, 2012) as the *silver* standard. Doing so enables us to compare the segmentation quality of a model trained on data from a single modality, in our case sMRI, with a model trained with the addition of connectivity data derived from dMRI. Furthermore, we explore the accuracy of different GNN architectures, as well as the influence of different sMRI-derived parameters on the final segmentation results.

We also validate the best-performing model by examining its effectiveness in a downstream task, thereby evaluating the applicability of the multimodal GNN and its possible improvement upon the silver-standard training set. Given that FreeSurfer segmentation labels are created automatically, as opposed to manually, they are not considered gold-standard ground truth (Fischl et al., 2004), and, as such, they might contain errors. Overlap measures between the labels resulting from GNN and those of FreeSurfer, therefore, cannot gauge any potential improvement over the training labels, as a perfect Dice score would only indicate that the model has learned to perfectly reproduce the FreeSurfer results. Conversely, disparities between the two results could indicate either an error on the part of the trained model or possibly an improvement upon the FreeSurfer segmentation, something that cannot be estimated with silver-standard labels alone. Evaluating the usefulness of the final segmentation when applied to a downstream task, such as age prediction, however, provides us with a surrogate measure to compare the two segmentation methods.

We describe our methods in Section 2, provide our experimental results in Section 3, and discuss them and conclude the paper in Section 4.

## 2 METHOD

The basis of our cortical graph structure is the pial surface automatically created with FreeSurfer (Fischl, 2012) from the T1-weighted 3D MRI image. The final result is in the form of a triangular mesh approximating the pial surface, which is shown in Fig. 1 for one of the test subjects. The resulting graph is unweighted. A feature vector is assigned to each vertex (node) in the graph and can contain a combination of the characteristics derived from sMRI and dMRI.

**Figure 1.**
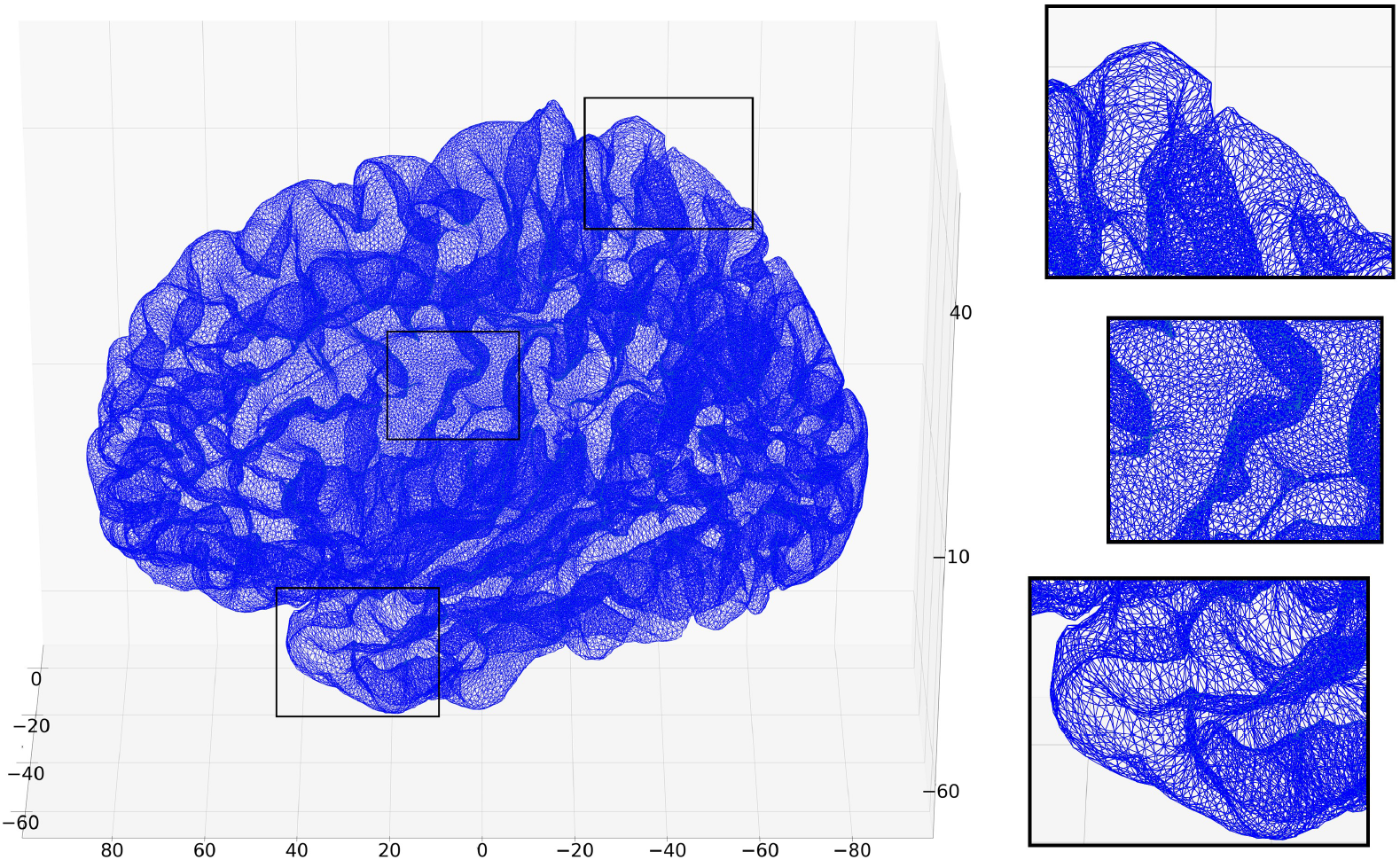
Visualization of the pial surface of an example subject from the training dataset. The zoomed-in regions show the triangular tessellation of the surface in greater detail. Note that while the structure of the graph is that of the mesh, the positioning of the nodes in the 3D space (used here for visualization) is not part of the graph. The (*x, y, z*) coordinates are in fact used as part of the feature vector of each vertex.

Data collected from sMRI that we incorporated in the feature vector include (*x, y, z*)-coordinates of the node, cortical thickness, and curvature values. Thickness is defined as the distance between the white and pial surfaces, and curvature is defined as the inverse of the radius of the circle tangent to the surface at the position of the corresponding vertex (mean curvature was used, which is the average of the curvatures corresponding to the smallest and the largest tangent circles). Thickness and curvature are scalar values, and the coordinates are a vector of length 3. The curvature is negative in a gyrus and positive in a sulcus. The features were min-max normalized.

To compute structural connectivity of each vertex to the other vertices from dMRI, we used a conductance-based method (Frau-Pascual et al., 2019). This approach models white-matter pathways within the brain based on a combination of differential Maxwell’s equations and Kirchoff’s circuit laws. A high conductance (i.e., low resistance) value computed between two regions represents a high degree of connectivity between them. This method accounts for both direct and indirect brain connections and has been shown to predict age and dementia (Frau-Pascual et al., 2021). We applied this method to sMRI and dMRI of 97 subjects of the “100 Unrelated Subjects” group of the Human Connectome Project (HCP) Young Adult study, which consists of healthy individuals aged 22-35 (Van Essen et al., 2013). Three subjects were omitted due to failed connectivity computation. We were granted access to restricted HCP data, which contains, among other sensitive information, the subjects’ exact age, Mini-Mental State Exam (MMSE) scores, and education level. During preprocessing, we transformed the data from the FreeSurfer uniform space, *fsaverage3*, where the brain connectivity data had been computed, to the native space of every subject via the *mris apply reg* command. The final connectivity map for each vertex was a feature vector of length 642 (the number of vertices in the hemisphere).

We used the following three different GNN architectures for segmentation:

- **Graph Convolutional Network (GCN)** is based on the Convolutional Neural Network concept, but designed to work on graph structures. With each layer, the information from the neighboring nodes is incorporated in the next layer’s feature vector (Kipf and Welling, 2017).
- **Graph Attention Network (GAT)** allows to assign a degree of usefulness to each neighboring node, making it a more complex architecture than the GCN (Veličković et al., 2018).
- **Graph U-Net** is based on the U-Net model, a deep-learning architecture well suited for the segmentation of biomedical images. It contains Graph Pooling layers, which down-sample the graph, followed by Graph Unpooling layers, which expand it back to the original size (Gao and Ji, 2019).

## 3 RESULTS

We first report the outcomes of the GNN model training on the task of cortical segmentation with the FreeSurfer segmentation as ground truth. Next, we detail the results of predicting the demographic/clinical characteristics of the subjects based on the segmentation results, which compares the usefulness of the model’s and the FreeSurfer’s outputs for a real-life task.

### 3.1 Graph-based cortical segmentation

All three models were evaluated on 97 subjects of the “100 Unrelated Subjects” group of the HCP Young Adult dataset. We held out 8 subjects as the final validation set, and the remaining 89 subjects were included in a 5-fold cross-validation. We monitored the training process in each split by examining the Dice scores of the model applied to both the training and the validation sets of the fold. In each fold, once training concluded, we applied the trained model to the final validation set (8 subjects). The mean Dice score on the final validation set over the five splits is reported and used as the primary metric to compare different models in the following.

For GNN model training, the Adam optimization algorithm and the *ReduceLRonPlateau* scheduler were used. The starting learning rate was *lr* = 0.01, and all models were trained for 100 epochs. Due to GPU constraints, a batch size of 1 was chosen. Gaussian noise was added to the input features during training for data augmentation.

Table 1 shows the final average Dice scores for the three model architectures and six different feature vectors, for the right and left hemispheres. For ease of reading, the model trained on coordinates, thickness, and curvature will be referred to as the “fully structural” model, and the model trained on all possible attributes, that is, coordinates, thickness, curvature, and structural connectivity, will be called the “multimodal” model.

**Table 1.**
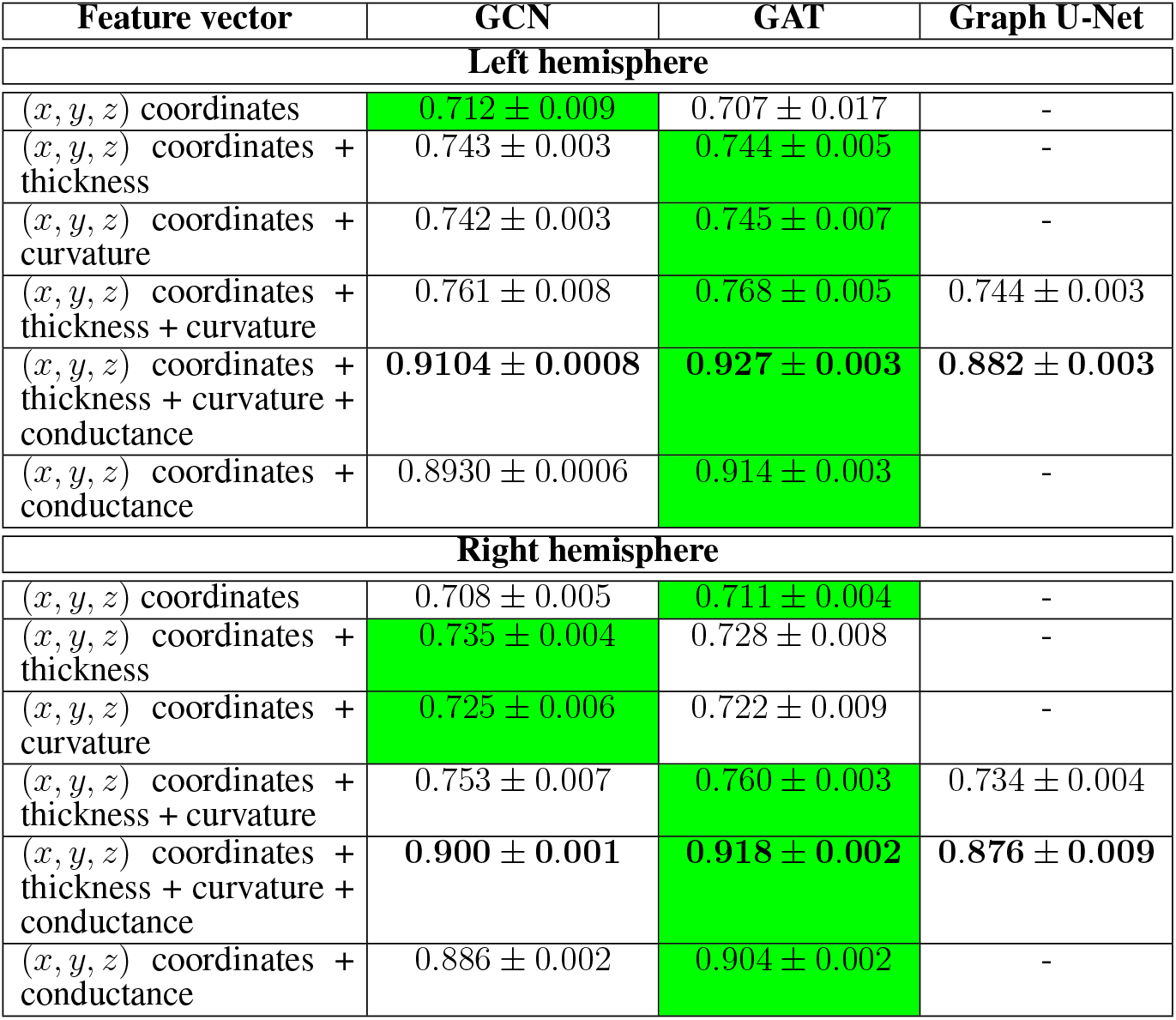
Dice scores (mean ± StD) on the final validation dataset for GCN, GAT, and Graph U-Net architectures for both left and right hemispheres. The highest value in each row is highlighted in green. The row with the highest values is in bold. Graph U-Net was trained only for two types of input feature vectors due to the significantly longer training time required and poor initial results.

#### 3.1.1 Influence of the Feature Vector

As can be seen in Table 1 for both hemispheres, using only (*x, y, z*) coordinates of the vertex produced the lowest Dice scores. Adding either the thickness or curvature scalar value slightly improved the results, with thickness often slightly more helpful than curvature. Adding them both further increased the final score, suggesting non-overlapping information by them. The most notable finding is the results by the multimodal model (when the connectivity values are added), where the Dice score is considerably increased, e.g. up to a cross-hemisphere mean of 0.923 for the GAT model. Worth pointing out is also that the model trained on the combination of (*x, y, z*) + conductance still outperformed all models trained on data solely from sMRI.

#### 3.1.2 Comparison of the model architectures

In terms of model performance, GAT outperformed GCN in nine out of the twelve feature combinations, including all that include connectivity values. Among models only trained on structural data, GCN either outperformed the GAT model or scored slightly below it. Graph U-Net achieved the lowest final scores in all examined cases.

Two-sided paired *t*-tests across the results on the final validation set revealed the subject-mean Dice scores from the multimodal model to be significantly higher for GAT as compared to GCN (*p* = 2 × 10^−5^ and *p* = 10^−6^ for the left and right hemispheres, respectively) and Graph U-Net (*p* = 7 × 10^−9^ and *p* = 3 × 10^−6^ for the left and right hemispheres, respectively).

#### 3.1.3 Influence of the number of parameters on the model performance

As detailed in Section 3.1.1, the addition of connectivity values to the feature vector considerably improved the segmentation results. For a fair comparison, the depth and width of the models were kept constant regardless of the size of the feature vector. However, as shown in Table 2, the connectivity data increase the size of the feature vector by 642, in turn increasing the number of trainable weights in our multimodal models, whereas thickness and curvature each only add a single scalar value per vector. In other words, when the sizes of the rest of the layers are kept constant, the number of trainable parameters will increase with the length of the input feature vector. For example, for the GCN architecture, the number of weights increases for a model trained solely on coordinates from 144k to 145k for a model trained on a combination of coordinates, thickness, and curvature, and finally up to 474k for a model trained on all available attributes.

**Table 2.** The influence of the number of model parameters on the final Dice score for a GCN model.

| $(x, y, z)$ coordinates + thickness + curvature | |
| --- | --- |
| Number of parameters | Dice Score |
| 145001 | 0.7410 ± 0.0013 |
| 285801 | 0.7633 ± 0.0031 |
| 499201 | 0.7630 ± 0.0017 |
| $(x, y, z)$ coordinates + thickness + curvature + conductance | |
| Number of parameters | Dice Score |
| 473705 | 0.9104 ± 0.0008 |

It is worth pointing out that due to the possibility of overfitting, increasing the number of trainable parameters does not always lead to enhanced performance. Having more parameters is therefore not necessarily an advantage for a model trained on the connectivity values. However, for the sake of completeness, we trained a GCN model with an increased width of the second layer on a feature vector solely including attributes from sMRI, to see how much it would affect its performance. As can be seen in Table 2, the final Dice score of the fully structural model is still significantly lower than the multimodal model, even with more trainable weights in the former than in the latter. When the number of features in the fully structural model is doubled relative to the original size, the score increases by 0.0223. However, upon increasing the size again, the result plateaus.

### 3.2 Downstream task

We further evaluated the performance of the GNN model by employing it in a downstream task, which tests the usefulness of the final segmentation results. As the multimodal GAT was the best-performing model in both hemispheres in Section 3.1.2, we chose this model for the downstream task.

#### 3.2.1 Statistical correlation

To examine the applicability of the segmentation outputs of the trained model and FreeSurfer, we correlated region-wise imaging attributes, namely average cortical thickness and average conductance, with demographic/clinical characteristics such as age, MMSE score, and education level. Specifically, we calculated the average thickness/conductance value for every segmented cortical region for both the FreeSurfer and the GAT model segmentations, and then determined how the resulting values correlated with the subjects’ age/MMSE/education.

When correlating the average of all within-region mean thickness values with age, for FreeSurfer segmentation, the left hemisphere showed statistical significance with *r* = −0.35 and 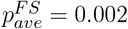. The negative linear correlation is consistent with what is reported in the literature (Aycheh et al., 2018). For the right hemisphere, the values are slightly less significant, with *r* = −0.23 and 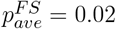. Model-based segmentation performed similarly to FreeSurfer-based segmentation; for the left hemisphere, *r* = −0.31 and 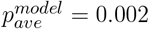, and for the right hemisphere *r* = −0.23 and 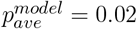.

In region-wise correlation, four FreeSurfer regions passed the Bonferroni-corrected threshold (*α*_*B*_ = 0.05*/*35 = 0.0014) in the left hemisphere and two in the right hemisphere. For model-based segmentation, five regions fulfilled the threshold for the left hemisphere, and two for the right hemisphere. Table 3 shows the *p*-values for the specific regions in which at least one of the segmentation sources passed the significance threshold *α*_*B*_. While this shows certain regions’ strong correlation with cortical thickness, the model and FreeSurfer performed similarly, as a paired Wilcoxon signed rank test on the region-wise *p*-values revealed no statistically significant difference (*p*_*W*_ = 0.16 and *p*_*W*_ = 0.88 for the left and right hemispheres, respectively).

**Table 3.** Brain regions with mean cortical thickness statistically significantly correlated with age (by at least one method), for FreeSurfer and GNN-based (GAT) segmentation.

| Region | FreeSurfer | GAT Model |
| --- | --- | --- |
| **Left Hemisphere** |  |  |
| Superior frontal gyrus | $p = 0.002,$<br>$r = -0.332$ | $p = 0.0008,$<br>$r = -0.334$ |
| Inferior frontal gyrus, Pars triangularis | $p = 1.86e - 07,$<br>$r = -0.500$ | $p = 9.3e - 07,$<br>$r = -0.474$ |
| Inferior parietal cortex | $p = 0.002,$<br>$r = -0.315$ | $p = 0.001,$<br>$r = -0.321$ |
| Superior temporal gyrus | $p = 0.001,$<br>$r = -0.321$ | $p = 0.0009,$<br>$r = -0.332$ |
| Banks of the superior temporal sulcus | $p = 7.0e - 05,$<br>$r = -0.392$ | $p = 7.7e - 05,$<br>$r = -0.390$ |
| **Right Hemisphere** |  |  |
| Middle frontal gyrus, Rostral division | $p = 0.0011,$<br>$r = -0.325$ | $p = 0.0009,$<br>$r = -0.332$ |
| Middle frontal gyrus, Caudal division | $p = 0.0001,$<br>$r = -0.383$ | $p = 0.0002,$<br>$r = -0.366$ |

No correlation between other brain attributes in a brain region and/or other demographic/clinical features passed the *α*_*B*_ significance threshold.

#### 3.2.2 Age prediction

For the prediction task, we combined regions from both hemispheres, resulting in an input vector of length 70. We used seven available machine learning models from the MATLAB Regression Learner Toolbox. We also trained different-sized Multi-Layer Perceptrons (MLPs). The performance was determined with the root mean square error (RMSE), which should be at the very least below the standard deviation of the label data, *σ*_*age*_ = 3.70 years, in order to show any form of utility.

FreeSurfer segmentation outperformed the GAT segmentation, achieving a lower RMSE for four out of seven models when using thickness as a brain characteristic and for all models when using conductance. However, the results for both methods were not satisfactory enough for a clear conclusion - the lowest reported RMSE was 3.44, i.e. only 0.26 below *σ*_*age*_, for the Regression Trees when using the FreeSurfer segmentation and the average conductance.

MLP showed poor training results, either indicating over- or underfitting, and a large variation in the RMSE between different validation splits. This significant training instability suggests that the model was not capable of properly generalizing on the dataset, making the comparison between the two segmentation methods difficult to interpret. As such, no strong conclusions could be drawn from the final results.

## 4 DISCUSSION

We demonstrated that GNNs are a suitable network architecture for the task of cortical surface segmentation, comparable to the non-deep-learning based algorithms such as the one employed by FreeSurfer. While the direct comparison with other methods in the literature is limited due to the difference in datasets and cortical atlases used, the final reported scores of 0.768 for the model solely trained on sMRI-derived data and, especially, 0.927 for the model trained by including connectomic data are comparable to the currently reported state-of-the-art results (Henschel et al., 2020, 2022; Roy et al., 2018; Wu et al., 2019; Cucurull et al., 2018; Eschenburg et al., 2021).

The considerable increase in the final Dice scores for the model trained in a multimodal fashion indicates the significant utility of structural connectivity in the task of brain segmentation. Notably, the model trained on the combination of (*x, y, z*) and conductance outperformed models trained on all available features from sMRI, which suggests that connectivity values provide more useful information than the thickness or curvature, even though those are the parameters that are utilized by the FreeSurfer segmentation algorithm (Fischl et al., 2004).

When considering the network architecture, GAT was the strongest one when connectivity data were included. However, GCN performed comparatively well when only sMRI values were considered. This is possibly because the more complicated architecture of the GAT model performs better when it needs to generalize on a more complex training set, and the GCN is not powerful enough to process the significantly longer feature vector once the connectivity data are introduced, even when it has more training weights. When the training data include only attributes from sMRI, a less powerful model such as a GCN might be enough.

It is worth noting that the Graph U-Net layers needed to be kept narrower than those in the other two architectures due to memory constraints, resulting in a much smaller number of training weights, 170403, for a Graph U-Net, in comparison to 473705 for a GCN and 242863 for a GAT model (for a feature vector of length 647). However, the fact that the GAT model, while having significantly fewer training weights than the GCN model, still managed to outperform it, indicates that a suitable model architecture should still be able to perform well with fewer trainable parameters. While GPU time and memory were limiting during model training, they are often much less of a concern during inference (e.g., while applying a trained model in a clinical setting), where even CPUs can be used to segment an image in a reasonable time.

The influence of trainable weights on the network performance was further explored in Section 3.1.3. The results from Table 2 indicate that, as expected, simply increasing the training weights is not a suitable alternative to introducing new attributes in the feature vector and adding new information to the training set.

In the second part of this work, we aimed to examine whether the GNN model has the capability to generalize upon FreeSurfer labels well enough to show higher utility in a downstream task posed. However, the results failed to unequivocally prove an improvement in the GNN-based segmentation. The RMSE from the MATLAB toolbox models did not show clear superiority of one method over the other, and the MLP showed typical behavior of an overfitted model, suggesting that the model most likely simply learned to memorize the input vector and their age labels, and did not have enough data to generalize on. It is therefore not possible to state whether the model was able to improve upon the silver-standard labels it was trained on.

One significant limitation of the study is the lack of manually labeled data, which could be used as the gold standard during training, serving as more reliable labels than the silver-standard segmentation created with FreeSurfer, and also as a benchmark for the evaluation of the final results. Training on more subjects could also improve the performance of the model in the downstream task. Nonetheless, as graph segmentation is intrinsically a node classification task, 97 subjects were enough for high training and validation scores. For the model to improve upon FreeSurfer, however, it may require a larger amount of data to be able to generalize upon the variation in the input. It is possible that with the amount of data provided, the model was capable of matching FreeSurfer’s segmentation, but not necessarily improving upon it.

Furthermore, it would be beneficial to see if the models are equally capable of generalizing on a more diverse group of subjects. As stated before, the brain surfaces used for training were all from a young and healthy population. However, for neuroscientific research as well as in clinical settings, patients and elderly individuals are often of more interest. Even more insightful would be to compare the influence of brain connectivity on the segmentation of the brains of patients with neurological disorders that are known to influence both the structure and connectivity of the brain, such as Alzheimer’s disease.

Future work includes validating our approach on gold-standard manually annotated cortical labels, attention-weight and node-importance analysis of the GAT architecture to explore which regions or connectivity features drive improvements, and training an end-to-end segmentation model to optimize downstream tasks (such as age prediction). Finally, more recent architectures, such as GCNs with an added attention mechanism (Tan et al., 2025) or diffusion-based approaches (Zhu et al., 2024), have shown great potential in the task of cortical surface segmentation. It would be worth exploring whether such state-of-the-art architectures could also benefit from the inclusion of structural connectivity.

## AUTHOR CONTRIBUTIONS

AŁ: Conceptualization, Formal analysis, Investigation, Methodology, Validation, Visualization, Writing – original draft, Writing – review & editing. AK: Conceptualization, Supervision, Writing – review & editing. CP: Conceptualization, Supervision, Resources, Writing – review & editing. AFP: Data curation, Resources, Writing – review & editing. IA: Conceptualization, Data curation, Funding acquisition, Project administration, Resources, Supervision, Writing – review & editing.

## DATA AVAILABILITY STATEMENT

Magnetic resonance images were provided by the Human Connectome Project (HCP) (Van Essen et al., 2013), WU-Minn Consortium (Principal Investigators: David Van Essen and Kamil Ugurbil): https://www.humanconnectome.org/study/hcp-young-adult.

The conductance model of structural brain connectivity (Frau-Pascual et al., 2019) was computed using our public toolbox at https://github.com/ainafp/FVT4DWI. FreeSurfer (Fischl, 2012) is publicly available at https://surfer.nmr.mgh.harvard.edu. The models employed here were built using the PyTorch Geometric library (Fey and Lenssen, 2019).

